## Supplementary figure 1-5 for "Cancer-associated hypersialylated MUC1 drives the differentiation of monocytes into macrophages with a pathogenic phenotype"

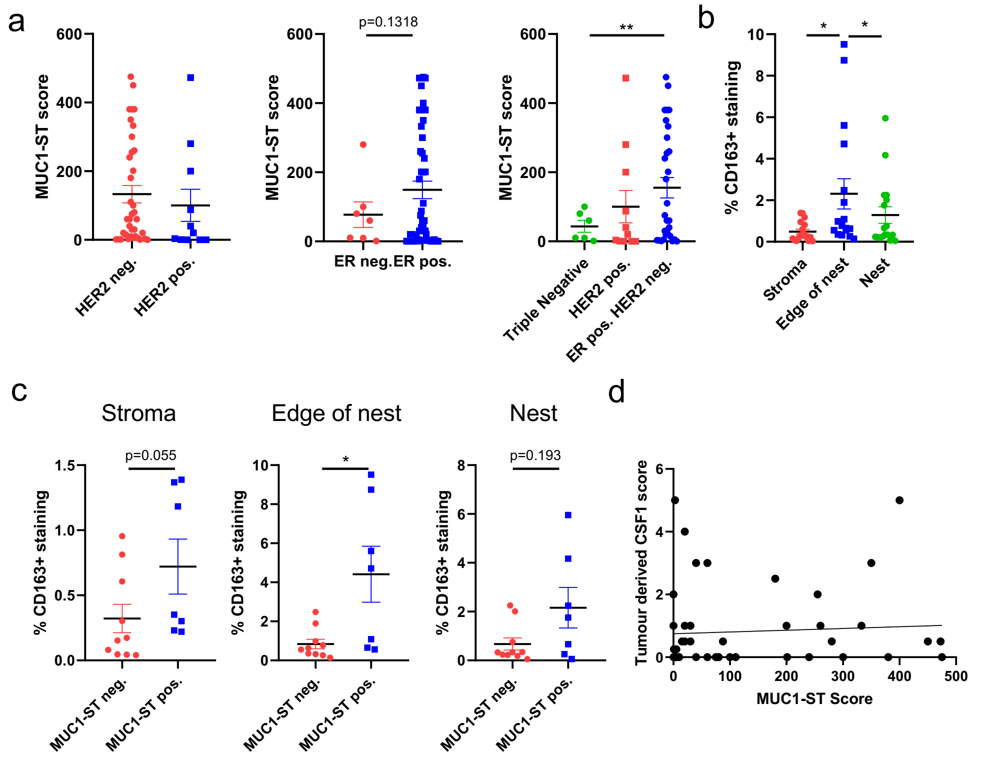


Figure S1. **(a)** Pathological subtyping in relation to MUC1-ST scoring (n=53). **(b)** % CD163 staining in different regions of tumours (n=17). **(c)** % CD163 staining in MUC1-ST positive (score >5) or negative (score <5) tumours in the indicated regions (n=17). **(d)** Correlation between MUC1-ST score and tumour-derived CSF1 score (n=53). Standard error of mean shown. (a-b) *p<0.05, **p<0.01 using unpaired t-test with Welch's correction owing to unequal population variance. (c) *p<0.05 Mann Whitney test (used as populations not Gaussian). Correlations were analysed using linear regression analysis (Pearson's).

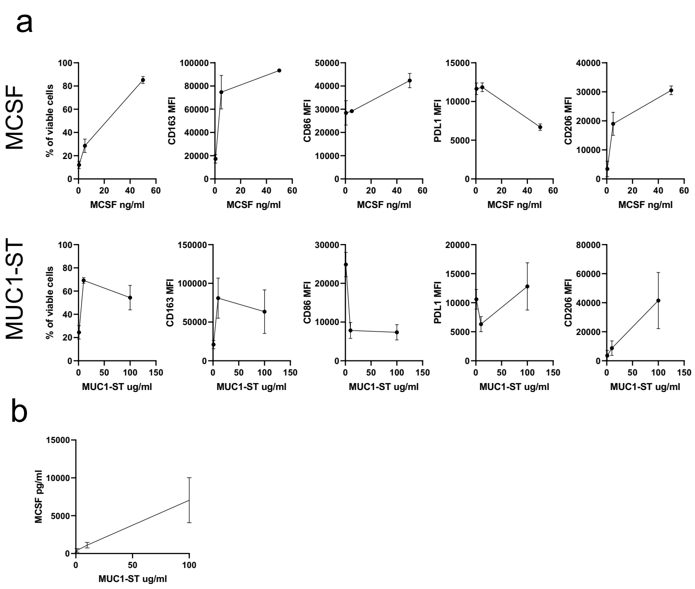


Figure S2. **(a)** The effect of different concentrations of MCSF (top row) and MUC1-ST (bottom row) on monocyte viability and phenotype (n=3). **(b)** The effect of different concentrations of MUC1-ST on MCSF release from monocytes at 48h (n=3). Standard error of mean shown.

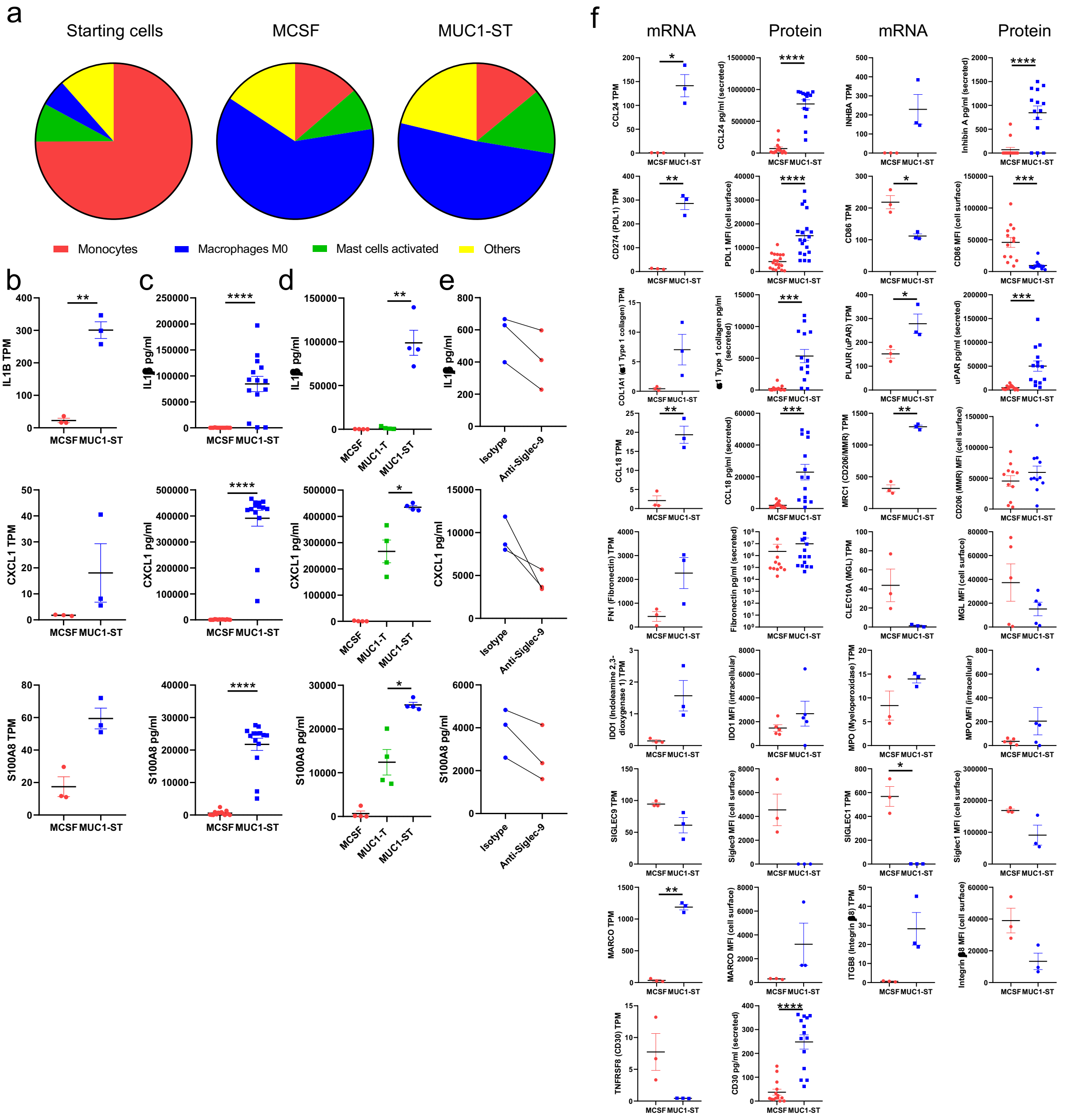


Figure S3 **(a)** Determination of immune cell subtype by applying CIBERSORT to starting monocytes (n=2) and monocytes treated with MCSF (n=3) or MUC1-ST (n=3) for 7 days **(b,c,d,e)** Three genes that are differentially expressed between MCSF macrophages and MUC1-ST macrophages **(b)** at the RNA level (n=3), **(c)** and at the protein level (n=14). **(d)** The same 3 proteins assessed in monocytes treated with desialylated MUC1-ST (MUC1-T) (n=4) **(e)** the effect of pre-incubation of monocytes with anti-Siglec-9 or isotype control, prior to MUC1-ST addition, on the production of these 3 proteins (n=3). **(f)** Assessment of an additional 17 proteins whose genes were differentially expressed between MCSF macrophages and MUC1-ST macrophages (n=3-14). Standard error of means shown. *p<0.05 **p<0.01 ***p<0.001, ****p<0.0001 using paired t-test. TPM; transcripts per million. MFI. Mean fluorescence intensity

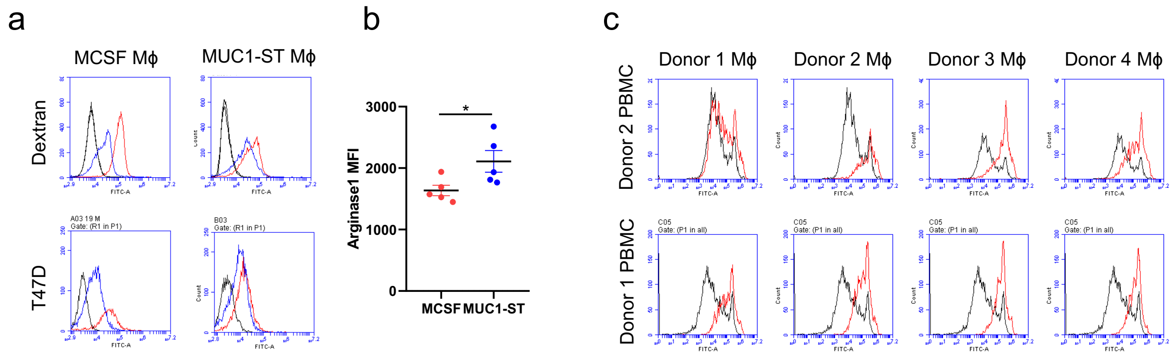


Figure S4. **(a)** representative histograms showing dextran-FITC uptake (top; blue; 4C. red; 37C) and the uptake of labelled T47D tumour cells (bottom; blue; 4C. red; 37C) by MCSF macrophages and MUC1-ST macrophages after 4h incubation, against macrophages alone (black). **(b)** intracellular staining of arginase in MCSF macrophages and MUC1-ST macrophages (n=5) **(c)** representative histograms showing the effect of media alone (black) or MUC1-ST (red) macrophage supernatant on the proliferation of CD3 stimulated PBMCs after 4 days. Standard error of mean shown, *p<0.05 using paired t-test.

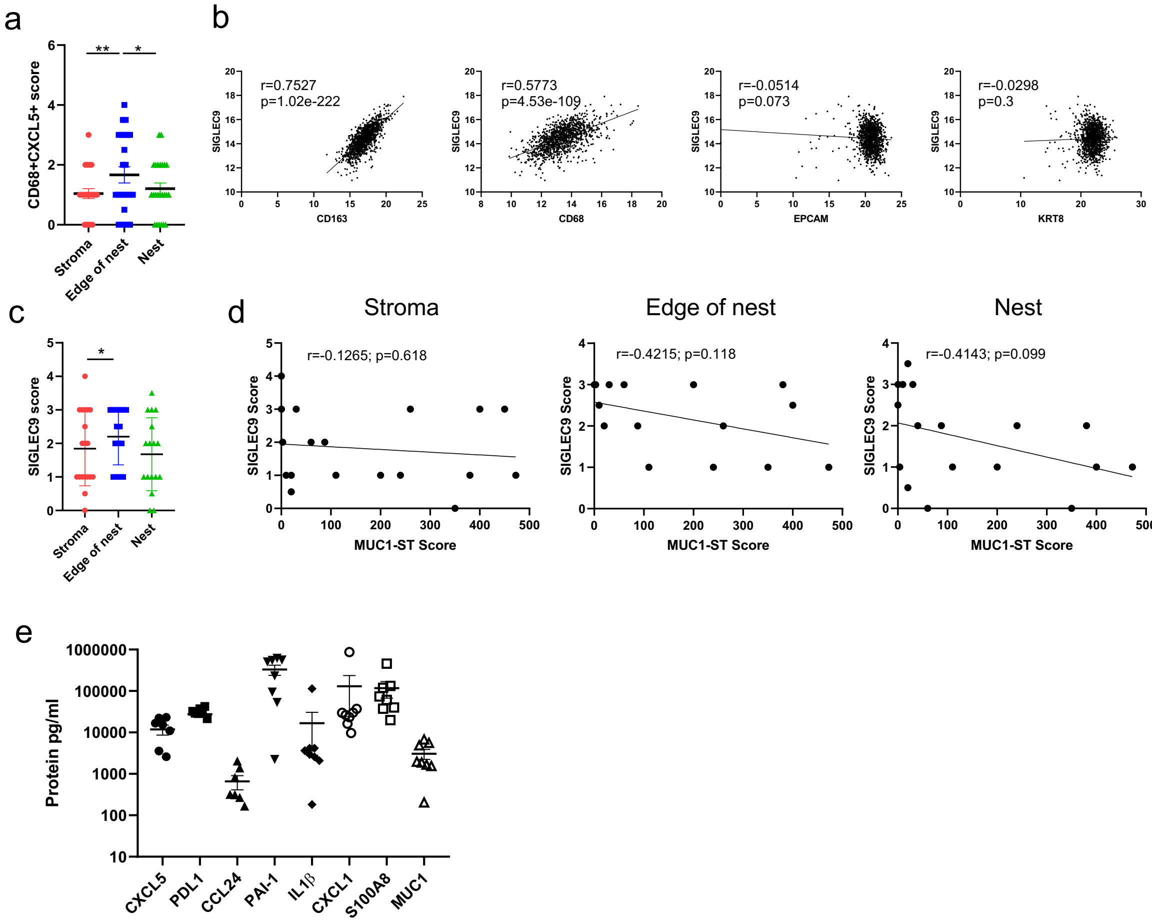


Figure S5. **(a)** Scoring of CD68+CXCL5+ cells in different indicated regions of the tumour (n=24). **(b)** TCGA data showing the correlation between SIGLEC9 and CD163, CD68, and EPCAM and KRT8 expression levels in breast cancers (n=1217). **(c)** Scoring of SIGLEC9+ cells in different indicated regions of the tumour (stroma n=18, edge n=15, nest n=17). **(d)** SIGLEC9 scores in different indicated regions of the tumour measured against MUC1-ST scoring (stroma n=18, edge n=15, nest n=17. **(e)** Measurement of indicated factors in the interstitial fluid of fresh breast cancers (n=8). Standard error of the mean shown and paired t-test used *p<0.05, **p<0.01. Correlations were analysed using linear regression analysis (Pearson's).
